# Projection layers improve deep learning models of regulatory DNA function

**DOI:** 10.1101/412734

**Authors:** Alex Hawkins-Hooker, Henry Kenlay, John Reid

## Abstract

With the increasing application of deep learning methods to the modelling of regulatory DNA sequences has come an interest in exploring what types of architecture are best suited to the domain. Networks designed to predict many functional characteristics of noncoding DNA in a multitask framework have to recognise a large number of motifs and as a result benefit from large numbers of convolutional filters in the first layer. The use of large first layers in turn motivates an exploration of strategies for addressing the sparsity of output and possibility for overfitting that result. To this end we propose the use of a dimensionality-reducing linear projection layer after the initial motif-recognising convolutions. In experiments with a reduced version of the DeepSEA dataset we find that inserting this layer in combination with dropout into convolutional and convolutional-recurrent architectures can improve predictive performance across a range of first layer sizes. We further validate our approach by incorporating the projection layer into a new convolutional-recurrent architecture which achieves state of the art performance on the full DeepSEA dataset. Analysis of the learned projection weights shows that the inclusion of this layer simplifies the network’s internal representation of the occurrence of motifs, notably by projecting features representing forward and reverse-complement motifs to similar positions in the lower dimensional feature space output by the layer.

## Introduction

The abundance of data characterising the function of non-coding DNA at high resolution facilitates the use of complex data-driven methods to learn the sequence features known as ‘motifs’ that encode this function (Alipanahi et al., 2015). A number of works have used neural networks to model human regulatory DNA, taking as input fixed-length regions of DNA sequence and predicting properties such as transcription factor binding, chromatin accessibility and histone marks using data collected by ENCODE and other consortia (Zhou and Troyanskaya, 2015; Kelley et al., 2016; Quang and Xie, 2016; Kelley and Reshef, 2017; Gupta and Rush, 2017; Zhou et al., 2018). Several of these networks are intended to simultaneously model a wide variety of the functional characteristics of the input region, by predicting hundreds or even thousands of such measurements across multiple cell types in a multi-task learning framework. With hundreds of known regulatory motifs recorded in databases such as JASPAR (Khan et al., 2018), machine learning models capable of fully characterising a significant variety of the measurable functional properties of human noncoding DNA must be able to recognise a large number of distinct patterns in the input sequence. Indeed while existing approaches have varied in the details of their neural network architectures, they have tended to share the use of relatively large numbers of convolutions as motif scanners in the first layer, and differed mainly in the subsequent layers where standard convolutions, dilated convolutions and recurrent layers have all been used to model interactions between features (Zhou and Troyanskaya, 2015; Kelley et al., 2016; Quang and Xie, 2016; Kelley and Reshef, 2017; Gupta and Rush, 2017). The best reported performance on the DeepSEA benchmark was achieved by a network having 1024 convolutional kernels in its first layer (Quang and Xie, 2016); indeed even when experimenting with single-output networks designed to predict binding for a single transcription factor, Zeng et al. (2016) observed that the performance of convolutional networks continued increasing with the number of filters in the first layer up to over 100 filters.^1^

The use of a sufficient number of first layer filters to capture the variety of motifs relevant to the task at hand thus appears to be an important consideration in the design of neural networks for processing noncoding DNA sequences. At the same time, it raises questions. For one thing, the use of a large number of parameters in the first layer raises the possibility of overfitting. Moreover, first layers designed to recognise large numbers of specific motifs are bound to produce outputs which are relatively sparse and high-dimensional, which may hamper learning in subsequent layers (Bengio et al., 2003). Finally, these layers are computationally expensive, particularly when applied to long sequences, both due to the cost of computing the activations by convolving the input at each point in the sequence, and the cost in the next layer of processing sequences of high-dimensional activation vectors.

Standard regularisation techniques such as dropout (Srivastava et al., 2014) may be expected to help alleviate the problem of overfitting, and have been applied to the first convolutional layer in previous works. But there is room for further work both in terms of characterising the extent of the problem and investigating alternative solutions. Projection layers, which can be used to reduce the dimensionality of a representation without reducing its resolution, are a popular component of deep networks in computer vision where they are often referred to as 1×1 convolutions (Szegedy et al., 2014; He et al., 2016; Lin et al., 2013). Reducing the dimensionality of a layer’s activations reduces the number of parameters required in the subsequent layer, as well as the cost of computing that layer’s activations. At the same time, depending on the nature of the features learned in the first layer, the denser representation resulting from the projection may well preserve much of the information contained therein. Even random projections are well known to preserve distances in dense representations (Johnson and Lindenstrauss, 1984; Bingham and Mannila, 2001).

The common practice of including amongst the training inputs both forward and reverse-complement versions of each target sequence in particular motivates the exploration of a more compressed representation. Models are forced by this form of data augmentation to recognise distinct instances (forward and reverse-complement) of functionally equivalent motifs. Methods capable of identifying these two instantiations therefore offer the same expressivity at potentially lower representational cost. Recognition of this issue has motivated the development of layers specially adapted to ensure the identity of forward and reverse-complementary sequences (Shrikumar et al., 2017). The use of projections offers an alternative approach to this problem.

Here we focus on the design choices related to the capacity of multitask networks to recognise a sufficient variety of motifs in input sequences, by jointly exploring both the effect of the number of first layer filters and the use of projection and dropout as approaches designed to mitigate the disadvantages of a large first layer. We choose to address these questions using the DeepSEA dataset (Zhou and Troyanskaya, 2015), since this has previously been used to benchmark different network architectures (Gupta and Rush, 2017; Quang and Xie, 2016). Initially using a reduced version of the dataset with shortened input regions, we vary the number of first layer filters for standard convolutional and convolutional-recurrent architectures with and without a projection layer and dropout, with our results indicating the importance of regularisation and the performance benefits of projection. We incorporate the projection layer into a convolutional recurrent neural network architecture with a number of modifications from the DanQ architecture proposed by Quang and Xie (2016). This new architecture achieves state of the art performance on the full DeepSEA dataset.

## Methods

### Baseline architectures

We experiment with modifications to two classes of architecture which have been successfully applied for multitask prediction in regulatory genomics. Details of the hyperparameters we used when training versions of these models are provided in the sections describing the relevant experiments.

1. CNN: Both DeepSEA (Zhou and Troyanskaya, 2015) and Basset (Kelley et al., 2016) use 3 layer CNNs, consisting of a stack of 3 convolution and max-pooling operations followed by one or more fully connected layers. DeepSEA’s convolutional layers are regularized using dropout and a global L2 penalty, whereas Basset applies batch normalization after each convolutional layer.
2. DanQ: The DanQ convolutional-recurrent architecture consists of a single convolutional layer followed by a pooling layer and a bidirectional LSTM (Graves and Schmidhuber, 2005). The full sequence of LSTM outputs are passed through two fully connected layers in order to generate predictions. Quang and Xie (2016) reported results for two versions of this architecture, DanQ and DanQ-JASPAR, differing in the sizes of the layers and in the initialization used for the first layer, with half of the better-perfoming DanQ-JASPAR’s 1024 first-layer filters being initialized using known motifs from the JASPAR database. Like DeepSEA, both DanQ architectures use dropout after their single convolutional layer.

### Linear projection layer

We investigate the use of a linear projection applied to the pooled activations of the first layer of architectures of both types. In detail, suppose that the first layer has *m* 1D convolutional filters and that after pooling the length of the sequence representation is *l*. Then the pooled activations form a sequence (*a*_1_, *a*_2_…*a*_*l*_) of m-dimensional vectors. The output of the projection layer is a sequence (*v*_1_, *v*_2_…*v*_*l*_) of *k*-dimensional vectors (*k* < *m*):

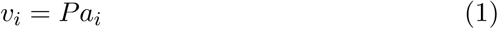

where *P* is a weight matrix of size *k* × *m*. The projection layer’s output is a sequence of the same length as the sequence of the first layer’s pooled filter activations, but whose members are vectors of a lower dimension, with the same projection matrix *P* being used to reduce the dimension at each point in the sequence. All the results reported below were obtained using a value of *k* = 64, which seemed to represent a good tradeoff between dimensionality reduction and preservation of information.

### Improved convolutional-recurrent architecture

The best previously reported performance on the DeepSEA dataset was achieved by the DanQ-JASPAR architecture which uses a single large convolutional layer followed by a max-pooling layer with stride and pool size of 15. This layer summarises the presence of the motifs identified by the convolutional layer across relatively large 15bp stretches of input. Pooling so aggressively has the advantage of controlling the length of sequence to be fed into the LSTM, preventing computation in the recurrent layer from becoming prohibitively time consuming. We hypothesise that this pooling involves throwing out useful positional information, which could be better preserved by splitting the downsampling across two sets of convolution and pooling layers rather than a single one. Therefore we propose an alternative convolutional recurrent (CRNN) architecture, which adds a projection layer, a second convolutional layer and a second pooling operation between the pooled outputs of the first convolutional layer and the bidirectional LSTM. To ensure fair comparison, the overall level of downsampling in the convolution and pooling layers is the same as in the DanQ-JASPAR networks, such that the length of the sequence of inputs to the bidirectional LSTM is the same (64) in both cases. In common with the DanQ networks we use a single fully-connected hidden layer before the output layer, but in order to control overfitting we use as input to this layer not the full sequence of LSTM outputs but their global mean. The proposed network, full details of which are given below, has far fewer parameters than DanQ-JASPAR and trains faster.

## Experiments

### The DeepSEA Dataset

The DeepSEA dataset consists of sequences of 1000bp from the human noncoding genome, labelled for the presence of a peak in the central 200bp in the signal for each of 919 chromatin features taken from ENCODE and Roadmap (Consortium, 2012; Consortium et al., 2015). These features represent a range of transcription factor binding, chromatin accessibility and histone modification measurements across a variety of cell types. Both forward and reverse-complement versions of the sequence corresponding to each set of targets are included in the dataset, meaning that models must be capable of learning both forward and reverse-complement motifs. We use the original training, validation and test splits and follow Quang and Xie (2016) in using as our primary evaluation metric test set AUPRC, which is calculated after averaging predictions across forward and reverse complement versions of each sequence.

### Design choices related to first layer on reduced DeepSEA dataset

In our first set of experiments we seek to rigorously explore the optimal configuration of the early layers of instantiations of both CNN and DanQ network designs. We vary the number of first layer filters, the use of dropout immediately after the first pooling layer, and the use of a projection layer (we fix the output dimension of this layer at each point in the sequence to 64) while keeping other hyperparameters fixed for a version of each class of architecture. When dropout and projection are used together, the dropout is applied after the projection layer. A dropout rate of 0.2 is used in all cases, which is the same as that applied to the activations of the first convolutional layer in both the DeepSEA and DanQ architectures. Other modifications to the original architectures were made in the interests of retaining comparable performance while reducing computational cost and are described below.

The CNN model that we choose to explore here takes from Basset the use of 3 convolutional layers, with kernel sizes of 19, 11 and 7 respectively, but varying in several other details. We use max pooling operations of sizes 6, 2, and 2 after the convolutional layers. The number of filters in the second and third convolutional layers is held fixed at 128 and 256 respectively. The outputs of the final pooling operation are fed into a single hidden layer of 2048 neurons to which dropout with dropout factor of 0.5 is applied. Leaky ReLUs (Maas et al., 2013) are used for all activations. Our DanQ architectures follow the original in most details other than those under investigation, except for the use of Leaky ReLU rather than ReLU activations, and the use of a reduced number of LSTM cells (100) in each direction.

To mitigate the cost of these experiments, we run them on a reduced version of the DeepSEA dataset, using only the central 500bp of each 1000bp sequence. For all networks we use the Adam optimizer (Kingma and Ba, 2014) with an initial learning rate of 3 × 10^−4^ to minimize the multitask binary cross entropy loss via mini-batch gradient descent with a batch size of 256. The learning rate was reduced by a factor of 5 if the validation loss did not decrease for two epochs. Training was terminated if the validation loss did not improve for five epochs. All models were implemented in Keras (Chollet et al., 2015) using the Theano backend (Theano Development Team, 2016).

### Evaluation of CRNN architecture on full DeepSEA dataset

For the second set of experiments we use the full 1000bp for each sequence and seek to compare the performance of our improved CRNN architecture to that of DeepSEA and the two DanQ architectures. For comparison with the two variants of DanQ, DanQ and DanQ-JASPAR, which have, respectively, 320 and 1024 filters in the first layer, we explore two variants of our CRNN architecture with 320 and 700 filters of length 30 in the first layer. To evaluate the contribution of the projection layer, for each CRNN variant we train one network with projection after the first pooling operation, and one network without projection but otherwise identical to the first. All networks use a second convolutional layer with 128 filters of length 11 whose activations are pooled and fed into a bidirectional LSTM with 300 units in each direction. Max-pooling with stride and pool size of 7 after the first convolutional layer and 2 after the second convolutional layer together with unpadded convolutions ensure that the sequence of inputs to the LSTM is of the same length as in the DanQ-JASPAR model. Dropout with a rate of 0.15 is applied to the projected first layer activations if projection is used, and to the pooled first layer activations if not. Recurrent dropout (Gal and Ghahramani, 2016) with a rate of 0.2 is applied to the LSTM. Leaky ReLUs are used for all layer activations. Networks are trained using the same learning rate schedules as in the previous set of experiments. We compare the average test set AUPRCs of our models with those of the publicly available trained DeepSEA and DanQ networks.

## Results

### Effects of first layer design choices on reduced-size dataset

In both fully convolutional and convolutional-recurrent architectures consistent benefits were achieved by increasing the number of first layer filters, with gradual saturation of performance (as measured by test set AUPRC averaged across the tasks) at around 1000 filters in both cases (Figure 1). In the fully convolutional networks the benefit of the projection layer was very clear, with all networks which used projection outperforming those that didn’t, often by considerable margins. A combination of dropout and projection achieved the best performance in every case. There is less evidence of benefit in the case of the networks using DanQ-style architectures, with networks with regularisation sometimes outperforming those without, but a lack of a clear pattern in the results, at least under the test set AUPRC metric. This is despite models incorporating dropout and the projection layer consistently achieving lower cross-entropy loss on the validation set. One factor in the difference between the two types of architectures is the degree of overfitting that the standard, unregularised architecture suffers. We observed that fully convolutional architectures showed a much greater tendency to overfit than convolutional-recurrent architectures (Figure 2). We note that unlike a convolutional layer, an LSTM already learns its own projection in the form of the weight matrix which transforms the inputs into the internal state space within the input and forget gates. These internal projections may help reduce both the tendency to overfit and the potential performance improvement associated with incorporating an additional projection layer. In contrast, inserting a projection layer into a CNN architecture substantially reduces the degree of overfitting (Figure 2), which allows CNN networks including projection layers continue to benefit from adding additional filters in the first layer, whereas without projection, CNN performance hardly improves beyond 500 first layer filters, as the benefit of extra feature detectors is offset by the increased likelihood of overfitting.

**Figure 1:**
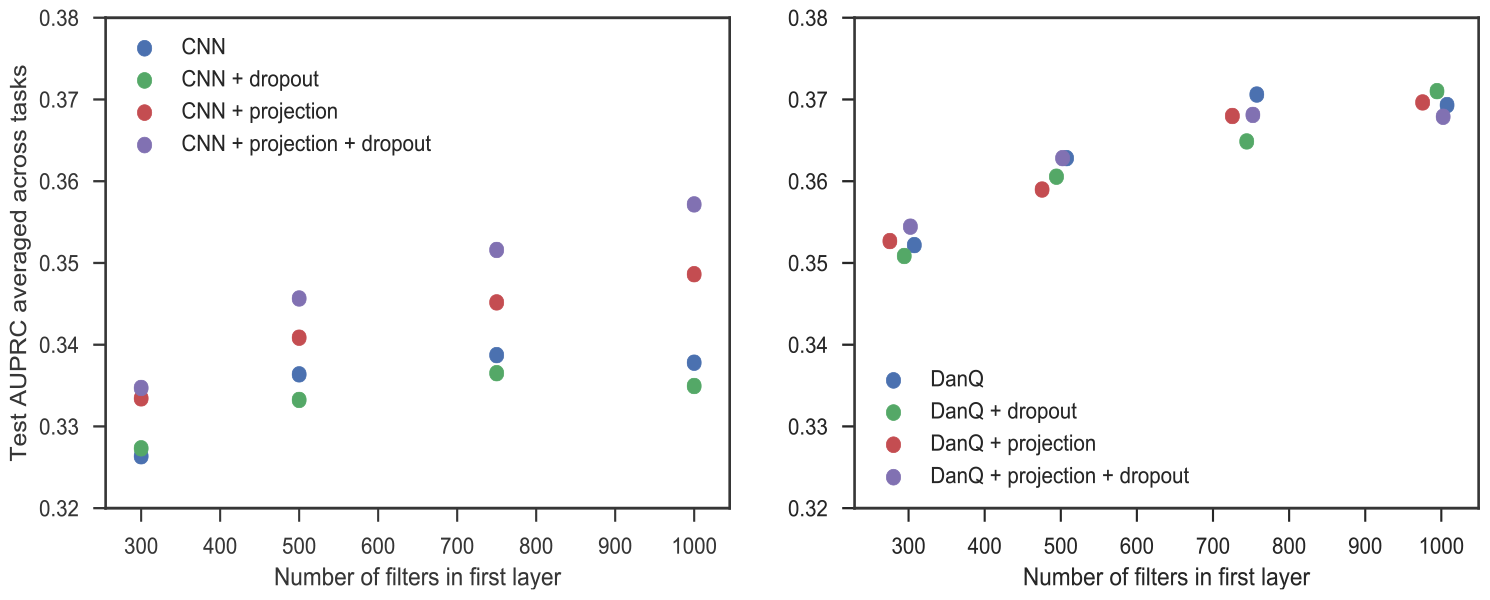
Test set AUPRC as function of first layer size for CNNs (left) and DanQ networks (right) with and without projection layer (projecting down to 64 dimensions) and dropout (dropout rate of 0.2). Jitter was added to the number of first layer filters for DanQ architectures to enable the points to be distinguished.

**Figure 2:**
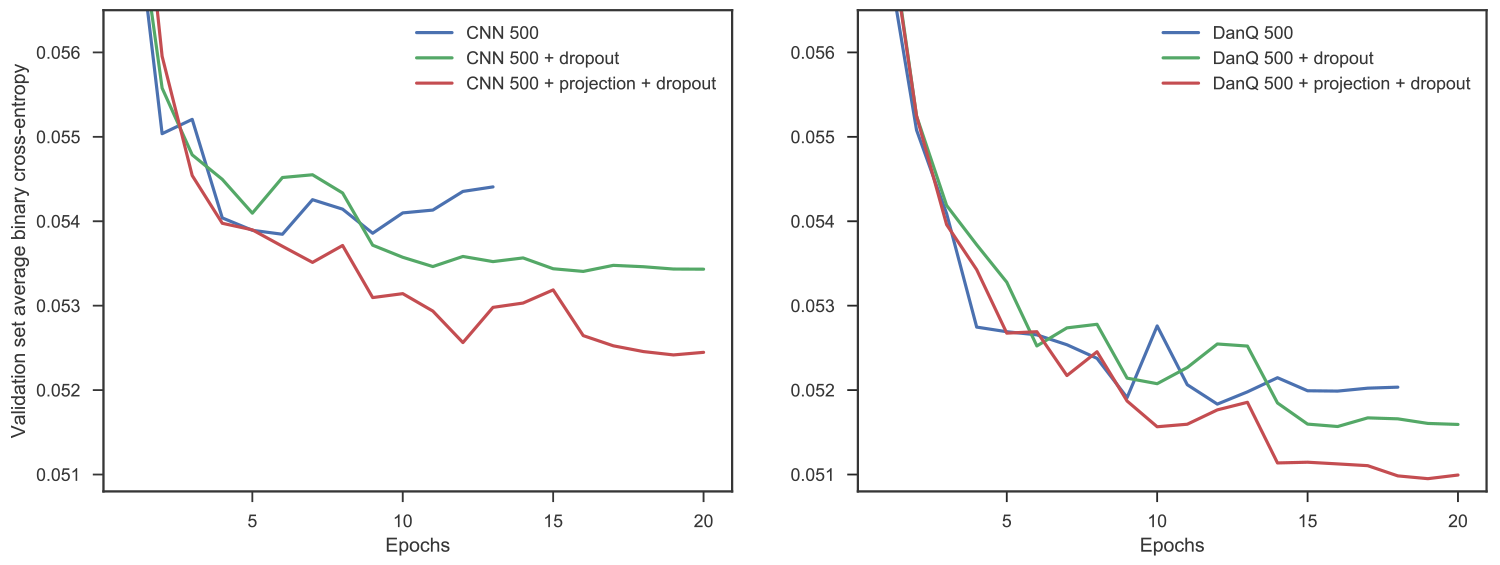
Validation set loss curves for CNN (left) and DanQ (right) models with 500 first layer filters, either with or without the two regularisation strategies. The CNN network shows much more evidence of overfitting.

### Projection layer helps improved CRNN architecture outperform other models on full DeepSEA data

Table 1 shows the cross entropy losses on the validation and test sets for our best-performing convolutional recurrent (CRNN) models as well as published baselines. CRNN-700 achieves the best average test set AUPRC of the compared models while being significantly less costly to train than DanQ-JASPAR, and without requiring the use of any known motifs to initialize first layer filters, as DanQ-JASPAR does. For both CRNN models we also compare the performance of models with and without the projection layer.

**Table 1:** Performance of CRNN models with and without projection layer compared to DeepSEA and DanQ networks. CRNN-n is a model with 2 convolutional layers with *n* and 128 filters respectively, with kernel sizes of 30 and 11, followed by a bidirectional LSTM with 300 units in each direction, whose outputs are averaged and fed through a hidden layer with 919 units which in turn feeds into the output layer. CRNN-*n*-projection is identical to CRNN-*n* except for the inclusion of a projection layer between the first and second convolution layers, which effectively reduces the dimension of the first layer’s activations from *n* to 64. Losses and AUPRCs for DanQ and DeepSEA networks are calculated using the publicly available model weights files. AUPRCs for all models are calculated after averaging predictions for forward and reverse complement versions of each test sequence, whereas forward and reverse complement versions of each sequence contribute independently to the reported losses.

| Model | Params | Valid Loss | Test Loss | Test AUPRC |
| --- | --- | --- | --- | --- |
| DeepSEA | 155,159,839 | 0.0509 | 0.0554 | 0.343 |
| DanQ | 46,926,479 | 0.0491 | 0.0538 | 0.371 |
| DanQ-JASPAR | 67,892,175 | 0.0482 | 0.0533 | 0.379 |
| CRNN-320-projection | 2,477,479 | 0.0485 | 0.0532 | 0.383 |
| CRNN-320 | 2,727,335 | 0.0489 | 0.0540 | 0.375 |
| CRNN-700-projection | 2,547,779 | **0.0475** | **0.0526** | **0.391** |
| CRNN-700 | 3,174,595 | 0.0484 | 0.0533 | 0.385 |

In both cases, the projection layer leads to a clear increase in performance and a reduction in the cost per epoch of training the network.

### Projection layer simplifies learning by unifying representations for forward and reverse-complement motifs

To understand the nature of the performance benefits brought by the use of the projection layer, we can investigate the relationship between the projection weights learned and the motifs learned by the first convolutional layer. To associate a motif with each filter in the first layer we follow a procedure similar to that introduced by (Alipanahi et al., 2015): several thousand sequences from the training set are passed through the trained model, and for each first layer convolutional filter we record the identities of the nucleotides at each position in the maximally-activating stretch of input in each sequence in which that filter is activated. From this we construct a PFM which can be converted into a motif representing the typical input pattern recognised by the filter. Using TOMTOM (Gupta et al., 2006) to search the JASPAR 2018 database (Khan et al., 2018) we find that 257 of the 700 learned motifs of the best-performing CRNN-700-projection model have at least one significant match (*q* < 0.01). Each learned motif is also associated with one of the columns in the 64 × 700 weight matrix of the projection layer. Suppose for example that at a certain point in an input sequence, the motif recognised by the *i^th^* convolutional filter occurs. Assuming none of the other filters are activated by this motif or its neighbouring region, the network’s representation of this region of the input will then just be the vector obtained by multiplying each weight in the *i^th^* column of the projection matrix by the filter’s activation. Thus the *i^th^* column of the projection matrix can be interpreted as representing an embedding of the motif learned by the *i^th^* convolutional filter. To visualise these embeddings, we choose to focus on a subset of the learned motifs which have the best matches to known motifs, selecting only the 44 learned motifs with *q*-values less than 10^−8^. The result of performing a PCA on the 44 columns of the projection weight matrix associated with these motifs is shown in Figure 3. Most strikingly, different versions of the same motif tend to cluster together, with the embeddings for filters which learn to recognise the forward version of a particular motif very often close to those for filters which recognise the reverse complement of the same motif. This suggests that the projection layer allows for a more efficient internal representation of motifs, recognising that forward and reverse complement patterns are functionally equivalent although completely different and therefore requiring different feature extractors at the sequence level. This representation of functional equivalence allows networks with a projection layer to harness the benefits of reverse-complement data augmentation without paying a price in terms of representational complexity.

**Figure 3:**
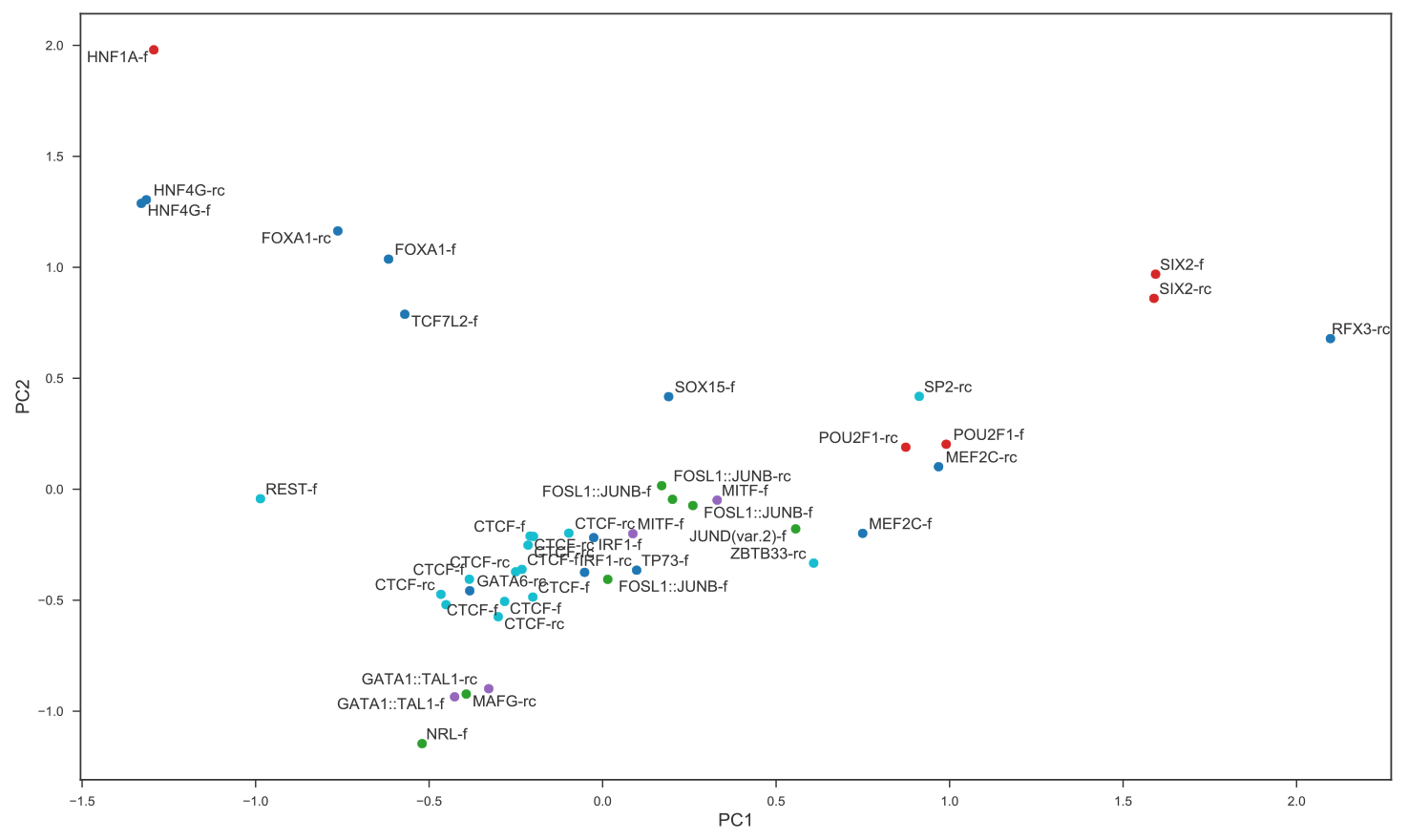
PCA of projection weights corresponding to learned motifs with best matches to known motifs in JASPAR database. Each point represents one of the 64 dimensional column vectors of the projection weight matrix. Only columns corresponding to learned motifs with a match with *q*-value less than 10^−8^ are included in the PCA to aid visualisation. Points are labelled by name of matched motif and whether it is the forward (f) or the reverse complement (rc) version of the known motif that is matched. Points are coloured by transcription factor family (cyan: C2H2 zinc finger, green: basic leucine zipper, red: homeodomain, purple: basic helix-loop-helix, blue: all other).

## Discussion

Despite the recent progress in the application of deep learning methods to model genomic data there remains work to be done in understanding the types of architecture and design choices best suited to the domain. We provide further evidence here that the performance of networks whose goal is to predict hundreds of functional properties from the DNA sequence is strongly dependent on the number of convolutional filters in the first layer. In networks where the subsequent layer is also convolutional, performance can be further improved by inserting a dimensionality-reducing projection layer between the two sets of convolutions. A similar use of projection layers in networks designed to predict enhancers was independently proposed by Chen et al. (2018) while we were finalising this manuscript. Their network takes as input both DNA sequences and chromatin accessibility information, and intersperses projections and convolutions on each of the two data modalities. While their work shows that projections can be used in highly performant architectures for regulatory genomics problems, they did not explore the role of projections in achieving this performance. Here our aim is to draw particular attention to the performance benefits and mode of functioning of a single projection layer, inserted directly after a first DNA motif-recognising convolutional layer, since we believe these point to its potential utility beyond any single application. In particular, we show that the projection layer is capable of learning the identity between forward and reverse-complement versions of functionally equivalent motifs and thereby simplifying the representation of the functional content of the sequence. It also reduces the number of parameters required in the subsequent layer, leading to less overfitting (particularly in combination with dropout) and reducing the computational cost. Incorporating the projection layer into a convolutional-recurrent network architecture similar to the DanQ architecture leads to improved performance on the DeepSEA dataset with fewer parameters and shorter per-epoch training times. Although we have only tested the use of the projection layer on the DeepSEA dataset, we believe that its use could be of important benefit in other situations in which accurate prediction of the targets requires recognition of a large variety of motifs in the input sequence.

## Contributions

AHH proposed the use of the projection layer, implemented and trained the models, ran the evaluations and wrote the paper. HK prototyped, implemented and trained early versions of the models and designed the evaluations. JR initiated the project, helped with the interpretation of the results and helped write the paper.

## Footnotes

1 For comparison, top performing networks on imagenet (He et al., 2016; Simonyan and Zisserman, 2014) make do with only 64 filters in the first layer despite the output dimension being comparable to that of DeepSEA. This discrepancy may perhaps be explained by the fact that while the features learned by the first layers of image processing networks are small, generic and only gradually composed into more specific features by subsequent layers, the features learned by the first layers of networks in regulatory genomics are by design highly specific motifs, typically 10-20bp in length.

## References

Alipanahi, B., Delong, A., Weirauch, M. T., and Frey, B. J. (2015). Predicting the sequence specificities of DNA-and RNA-binding proteins by deep learning. Nature biotechnology, 33(8):831–838.

Bengio, Y., Ducharme, R., Vincent, P., and Janvin, C. (2003). A neural probabilistic language model. J. Mach. Learn. Res., 3:1137–1155.

Bingham, E. and Mannila, H. (2001). Random projection in dimensionality reduction: Applications to image and text data. In Proceedings of the Seventh ACM SIGKDD International Conference on Knowledge Discovery and Data Mining, KDD ‘01, pages 245–250, New York, NY, USA. ACM.

Chen, S., Gan, M., Lv, H., and Jiang, R. (2018). DeepCAPE: a deep convolutional neural network for the accurate prediction of enhancers. bioRxiv.

Chollet, F. et al. (2015). Keras. https://github.com/fchollet/keras.

Consortium, R. E., Kundaje, A., Meuleman, W., Ernst, J., Bilenky, M., Yen, A., et al. (2015). Integrative analysis of 111 reference human epigenomes. Nature, 518(7539):317–330.

Consortium, T. E. P. (2012). An integrated encyclopedia of DNA elements in the human genome. Nature, 489(7414):57–74.

Gal, Y. and Ghahramani, Z. (2016). A theoretically grounded application of dropout in recurrent neural networks. In Lee, D. D., Sugiyama, M., Luxburg, U. V., Guyon, I., and Garnett, R., editors, Advances in Neural Information Processing Systems 29, pages 1019–1027. Curran Associates, Inc.

Graves, A. and Schmidhuber, J. (2005). Framewise phoneme classification with bidirectional lstm and other neural network architectures. NEURAL NETWORKS, pages 5–6.

Gupta, A. and Rush, A. M. (2017). Dilated convolutions for modeling longdistance genomic dependencies. ArXiv e-prints.

Gupta, S., Stamatoyannopoulos, J. A., Bailey, T. L., and Noble, W. S. (2006). Quantifying similarity between motifs. Genome Biology, 8.

He, K., Zhang, X., Ren, S., and Sun, J. (2016). Deep residual learning for image recognition. In 2016 IEEE Conference on Computer Vision and Pattern Recognition, CVPR 2016, Las Vegas, NV, USA, June 27–30, 2016, pages 770–778.

Johnson, W. and Lindenstrauss, J. (1984). Extensions of Lipschitz mappings into a Hilbert space. In Conference in modern analysis and probability (New Haven, Conn., 1982), volume 26 of Contemporary Mathematics, pages 189–206. American Mathematical Society.

Kelley, D. R. and Reshef, Y. A. (2017). Sequential regulatory activity prediction across chromosomes with convolutional neural networks. bioRxiv.

Kelley, D. R., Snoek, J., and Rinn, J. L. (2016). Basset: Learning the regulatory code of the accessible genome with deep convolutional neural networks. 26(7):990–999.

Khan, A., Fornes, O., Stigliani, A., Gheorghe, M., Castro-Mondragon, J. A., et al. (2018). JASPAR 2018: update of the open-access database of transcription factor binding profiles and its web framework. Nucleic Acids Res, 46.

Kingma, D. P. and Ba, J. (2014). Adam: A method for stochastic optimization. CoRR, abs/1412.6980.

Lin, M., Chen, Q., and Yan, S. (2013). Network in network. CoRR, abs/1312.4400.

Maas, A. L., Hannun, A. Y., and Ng, A. Y. (2013). Rectifier nonlinearities improve neural network acoustic models. In in ICML Workshop on Deep Learning for Audio, Speech and Language Processing.

Quang, D. and Xie, X. (2016). DanQ: A hybrid convolutional and recurrent deep neural network for quantifying the function of DNA sequences. 44(11):e107.

Shrikumar, A., Greenside, P., and Kundaje, A. (2017). Reverse-complement parameter sharing improves deep learning models for genomics. bioRxiv.

Simonyan, K. and Zisserman, A. (2014). Very deep convolutional networks for large-scale image recognition. CoRR, abs/1409.1556.

Srivastava, N., Hinton, G., Krizhevsky, A., Sutskever, I., and Salakhutdinov, R. (2014). Dropout: A simple way to prevent neural networks from overfitting. J. Mach. Learn. Res., 15(1):1929–1958.

Szegedy, C., Liu, W., Jia, Y., Sermanet, P., Reed, S. E., Anguelov, D., Erhan, D., Vanhoucke, V., and Rabinovich, A. (2014). Going deeper with convolutions. CoRR, abs/1409.4842.

Theano Development Team(2016). Theano: A Python framework for fast computation of mathematical expressions. arXiv e-prints,abs/1605.02688.

Zeng, H., Edwards, M. D., Liu, G., and Gifford, D. K. (2016). Convolutional neural network architectures for predicting DNA protein binding. Bioinformatics, 32(12) :i121–i127.

Zhou, J., Theesfeld, C. L., Yao, K., Chen, K. M., Wong, A. K., and Troyanskaya, O. G. (2018). Deep learning sequence-based ab initio prediction of variant effects on expression and disease risk. Nature Genetics, 50(8):1171–1179.

Zhou, J. and Troyanskaya, O. G. (2015). Predicting effects of noncoding variants with deep learning-based sequence model. Nature methods, 12(10):931–934.

